# Identifying content-invariant neural signatures of perceptual vividness

**DOI:** 10.1101/2022.11.30.518510

**Authors:** Benjy Barnett, Lau M. Andersen, Stephen M. Fleming, Nadine Dijkstra

**Author notes:** Joint senior authors (Stephen M. Fleming, Nadine Dijkstra).

## Abstract

Some conscious experiences are more vivid than others. Although perceptual vividness is a key component of human consciousness, how variation in this magnitude property is registered by the human brain is unknown. A striking feature of magnitudes in other psychological domains, such as number or reward, is the existence of neural magnitude codes that are independent from sensory contents. To test whether perceptual vividness similarly covaries with neural codes that are invariant to sensory content, we reanalysed existing MEG and fMRI data from two distinct studies, quantifying perceptual vividness via subjective ratings of awareness and visibility. Using representational similarity and decoding analyses, we find evidence for the existence of content-invariant neural codes for perceptual vividness distributed across visual, parietal, and frontal cortices. Our findings are consistent with a hypothesis that the subjective vividness of conscious experience is supported by neural signatures similar to magnitude codes in other cognitive domains.

**Significance Statement:** The vividness of conscious experience varies across different stimuli and contexts. Despite being a fundamental feature of conscious awareness, exactly how perceptual vividness is encoded in the human brain remains unclear. Neural codes underpinning magnitude in reward and numerosity domains have been shown to be unchanging as stimulus identity varies. Here, we test whether components of neural activity covarying with the magnitude of perceptual vividness are similarly independent of perceptual content in analyses of MEG and fMRI data. We find dynamic, content-invariant neural signatures of vividness in visual, parietal, and frontal cortices. Our findings introduce the surprising notion that neural signatures of conscious experience might follow similar coding principles to those found for magnitude properties of entirely different cognitive domains.

## Introduction

Humans are aware of their environment: we experience the sights and sounds of the world around us. Some experiences, however, are more vivid than others. For example, seeing a bird on a clear day will be more vivid than seeing one on a foggy evening. Similarly, a car alarm outside your office can be very vivid until your attention is consumed by a work task. The neural correlates of experience therefore naturally involve some representation of the magnitude of perceptual vividness. While the neural basis of perceptual vividness is yet to be systematically characterised, neural codes for magnitude quantities in other cognitive domains, such as reward and numerosity, are better understood. Many neural magnitude codes contain a content-invariant component, where the magnitude property is represented independently of its sensory features (Chib et al., 2009; McNamee et al., 2013; Piazza et al., 2007). For instance, the number “9” is represented as larger than the number “5”, regardless of whether we are comparing 9 vs. 5 apples, oranges, or saxophones. Here, we ask whether similar magnitude properties of perceptual vividness are also invariant to stimulus content: i.e. is the difference in vividness between seeing a bird on a clear day versus on a foggy evening represented in a similar way as the difference in vividness between hearing a car alarm with or without attending to it? We investigate this question by testing the extent to which neural signatures associated with reports of perceptual vividness are independent of perceptual content.

Content-invariance is a well-established feature of several neural magnitude codes. In the orbitofrontal cortex, for example, common representations of reward magnitude are shared across vastly different reward identities (Chib et al., 2009; Howard et al., 2015; Klein-Flügge et al., 2013; McNamee et al., 2013; Padoa-Schioppa & Assad, 2006). Furthermore, presentation of the same numerosity elicits suppression effects across symbolic (Arabic numerals) and non-symbolic (dots) stimuli in the intraparietal lobe (Piazza et al., 2007), and multivariate cross-classification has revealed common representations of numerosity across symbolic and non-symbolic formats too (Teichmann et al., 2018). Finally, in strong support for the content-invariance of magnitude codes, there is also evidence that numerical and reward magnitudes (amongst others) are encoded in a domain-general manner where, for example, higher numbers are represented similarly to highly rewarding stimuli and lower numbers are represented similarly to stimuli with low reward values, indicating a shared neural system underpinning representations of magnitude in both domains (Luyckx et al., 2019; Summerfield et al., 2020; Walsh, 2003).

Given the evidence for the existence of content-invariant neural magnitude codes in other domains, it is intriguing to ask whether invariance to perceptual content is also a feature of the neural activity covarying with the magnitude of perceptual vividness. If perceptual vividness is only encoded in a content-specific manner, our experience of a stimulus such as a red circle may become vivid through the increased firing of neural populations representing this feature (Itti & Baldi, 2009) (Figure 1, left). However, if neural activity covarying with perceptual vividness also contains a content-invariant component, we should be able to find neural signatures of vividness that are independent of those covarying with sensory features. The drivers of such content-invariant signals may include changes in attention, emotion, and other cognitive factors that surpass stimulus-specific salience, but nevertheless contribute to the vividness of experience (Morales, 2018, 2021) – an idea consistent with philosophical positions that distinguish percepts from their ‘force’ and ‘vivacity’ (Hume, 2000; Teng, 2022). As such, rather than being bound to content-specific representations, perceptual vividness might also covary with neural activity in a domain-general fashion, independently of stimulus content (Figure 1, right) (Levinson et al., 2021; Podvalny et al., 2019; Samaha et al., 2017; Sanchez et al., 2020).

**Figure 1.**
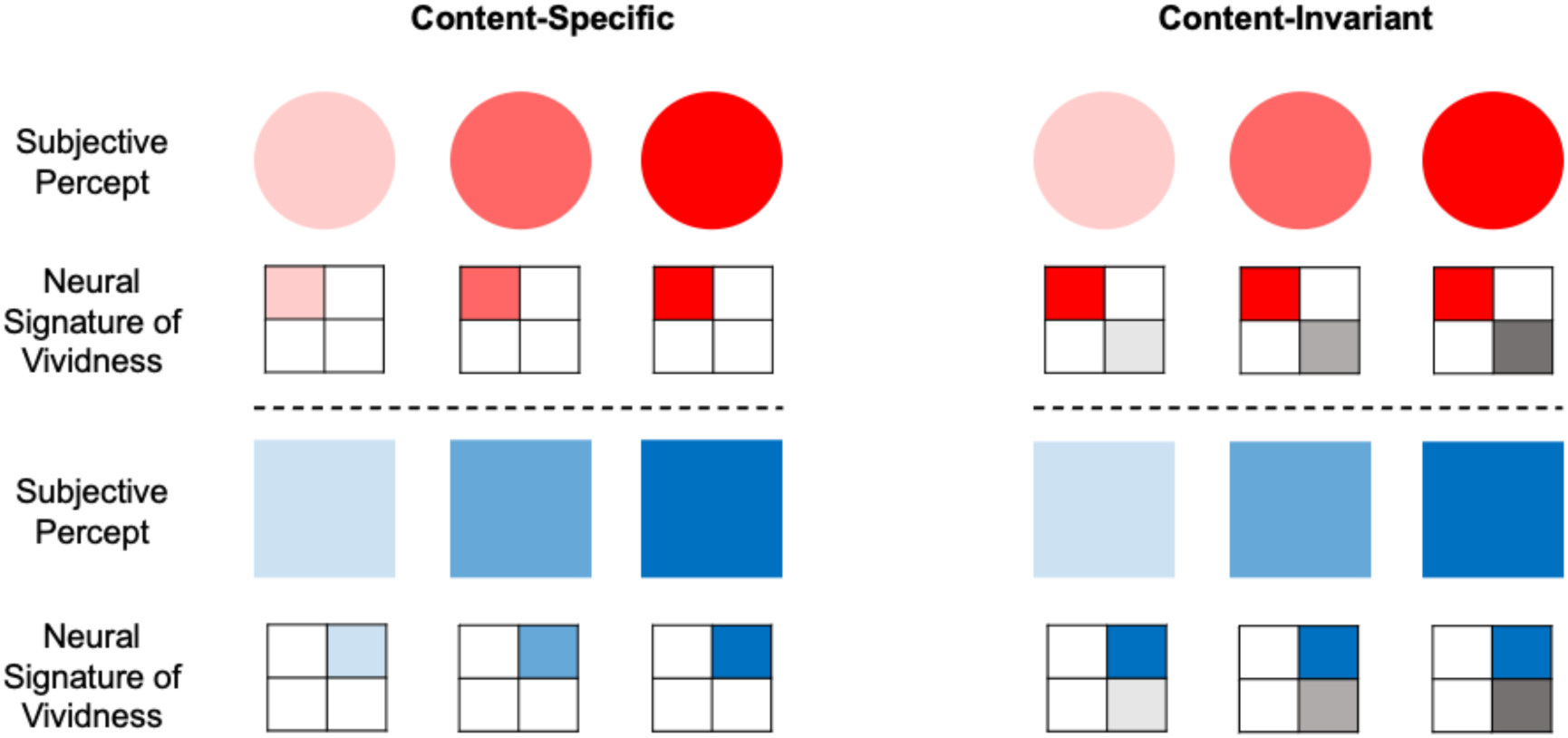
Hypothesised neural signatures of perceptual vividness. *Left: Content-specific neural signatures associated with perceptual vividness.* The subjective vividness of a red circle is associated with the strength of red circle-representing neurons while the vividness of a blue square is associated with the strength of blue square-representing neurons. The neural signatures underpinning the vividness of these two experiences are therefore different. *Right: Content-invariant neural signatures associated with perceptual vividness.* The subjective vividness of both red circles and blue squares is associated with the same neural signature, which supersedes the stimulus-specific neural activity. Attention, emotion, and other cognitive factors may drive this content-invariant neural signal of vividness. We note that the represented coding schemes are not mutually exclusive, and it is likely that both play a role in determining the vividness of perceptual experience.

To test whether the neural code for perceptual vividness contains a content-invariant component, we reanalysed both magnetoencephalography (MEG; Andersen et al., 2016) and functional magnetic resonance imaging (fMRI; Dijkstra et al., 2021) data to investigate how perceptual vividness is represented in the human brain. The difficulty in isolating pure correlates of vividness and awareness from co-varying neural signals (e.g. those related to arousal or performance) is well-known, and we did not attempt to tackle these issues here (Aru et al., 2012; Lau, 2022). Instead, we sought to determine the representational structure of awareness and visibility reports about different stimulus contents, to ask whether neural signatures covarying with vividness did so in a content-specific or content-invariant manner. To anticipate our results, we found evidence that neural representations of perceptual vividness generalize over stimulus content, exhibit a graded structure, and can be identified across visual, parietal, and frontal brain regions, consistent with signatures of magnitude codes in other cognitive domains.

## Materials and Methods

### MEG experiment

To explore the structure and dynamics of abstract representations of awareness ratings, we re-analysed an MEG dataset previously acquired at Aarhus University (Andersen et al. 2016). The data were recorded in a magnetically shielded room using an Elekta Neuromag Triux system with 102 magnetometers and 204 orthogonal planar gradiometers. Data were recorded at a frequency of 1000 Hz.

#### Participants

Nineteen participants took part in the experiment (Mean age = 26.6 years; SD = 4.4 years). Two participants were excluded from our analyses: one for failing to complete the experiment and the other for not using the ‘almost clear experience’ rating at all (see procedure details below).

#### Experimental Design and Statistical Analyses

In order to obtain a range of awareness ratings from each subject, a visual masking paradigm was used (Figure 2A). First, a fixation cross was presented for either 500, 1000, or 1500 ms, followed by a target stimulus for 33.3 ms. The target stimulus was either a square or a diamond presented in white/grey on a black (RGB value 0, 0, 0) background (Figure 2B). A static random noise mask followed the target and was presented for 2000 ms. Participants were required to identify the target during these 2000 ms, before rating their awareness of the stimulus on the perceptual awareness scale (PAS). PAS consists of four possible responses: no experience (NE), weak glimpse (WG), almost clear experience (ACE), and clear experience (CE). Following identification of the target, participants reported their awareness of the stimulus. The response boxes used for target identification and awareness report were swapped between hands every 36 trials to minimise lateralised motor responses contributing to MEG activity patterns. More details regarding the instructions given to participants about each PAS response can be found in Andersen et al. (2016).

**Figure 2.**
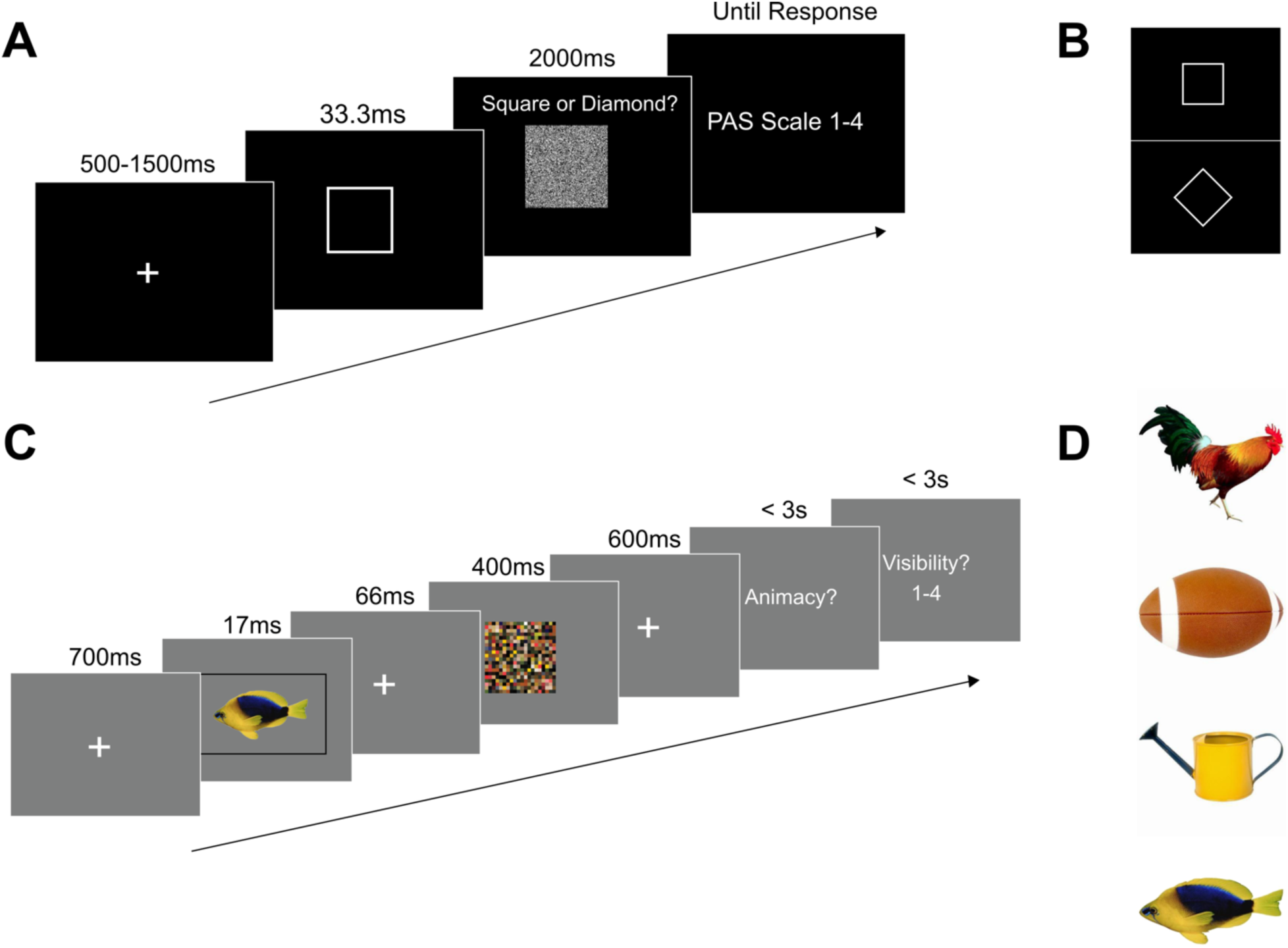
Experimental paradigms. **A:** Experimental paradigm for the MEG data collected by Andersen et al. (2016). First, a fixation cross was presented for 500, 1000 or 1500 ms. Then, either a square or a diamond was shown for 33.3 ms, followed by a static noise mask for 2000 ms. While the mask was shown, participants reported the identity of the target. Finally, they reported their awareness of the stimulus using the PAS scale. **B:** Stimuli used in Andersen et al. (2016). **C**: Experimental paradigm for the fMRI data collected by Dijkstra et al. (2021). A stimulus was presented for 17 ms, followed by a 66 ms ISI and a 400 ms mask. Participants then indicated whether the stimulus was animate or inanimate, and finally rated the visibility of the stimulus on a four-point scale. **D:** Stimuli used in Dijkstra et al. (2021).

The experiment consisted of one practice block and 11 experimental blocks, each with 72 trials. A contrast staircase was used for the target stimuli in order to obtain a sufficient number of responses for each PAS rating. The staircase procedure had 26 contrast levels ranging from a contrast of 2% to 77%, with a step size of 3 percentage points. In the practice block and first experimental block, the staircase increased by 1 level if a participant made an incorrect judgement on the identification task, and decreased by 2 levels if a participant made 2 successive correct identification judgements. For the remainder of the blocks, the staircase was adjusted based on which PAS rating the participant had used least throughout the experiment so far. Specifically, if NE had been used the least number of times throughout a block, 3 levels were subtracted after 2 consecutive correct answers, and 1 added for a wrong answer. If WG was the least used response, 2 levels were subtracted and 1 added. For ACE, 1 level was subtracted and 2 added. For CE, 1 level was subtracted and 3 added. This staircase procedure ensured a sufficient number of responses for each rating.

#### Pre-processing

MEG data were analysed using MATLAB 2019a and FieldTrip (Oostenveld et al., 2011). The data were pre-processed with a low pass filter at 100 Hz, as well as a Discrete Fourier Transform (DFT) and bandstop filters at 50 Hz and its harmonics. The data were split into epochs of −200 to 2000 ms around stimulus onset and down-sampled to 250 Hz. For baseline correction, for each trial, activity 200 ms prior to stimulus presentation was averaged per channel and subtracted from the entire epoch. During artefact rejection, trials with high variance were visually inspected and removed if they were judged to contain excessive artefacts. This procedure was performed blind to the experimental condition to avoid experimenter bias and was completed separately for the magnetometers and gradiometers in the Elekta Neuromag Triux system. Following artefact rejection the mean number of trials per PAS rating were as follows (numbers in brackets refer to standard deviations): No Experience: 180.35 (59.64); Weak Glimpse: 168.10 (74.75); Almost Clear Experience: 186.35 (82.74); Clear Experience: 115.24 (77.13). To further remove eye-movement artefacts, an independent components analysis was carried out on the MEG data, and the components with the highest correlation with each of the electro-oculographic (EOG) signals were discarded after visual inspection. Components showing topographic and temporal signatures typically associated with heart rate artefacts were also removed by eye.

Since we re-analysed previously collected data, we were unable to fully control for neural signals that typically covary with awareness level. As such, to better characterise the contribution of these signals to ratings of awareness we created two additional analysis pipelines. First, to ensure our results were not entirely driven by the contrast level of the stimuli, we regressed stimulus contrast level on each trial out of the pre-processed MEG data. Second, to investigate whether differences in pre-stimulus activity contributed to differences in perceptual visibility (Benwell et al., 2017; Podvalny et al., 2019; Samaha et al., 2017) we ran our data through the same pre-processing pipeline as above except for two adjustments: removing the baseline correction stage and lengthening the epochs to −450 ms to 2000 ms around stimulus presentation. The omission of baseline correction allows the analysis to be sensitive to differences in the pre-stimulus activity (in the offset or mean amplitude, for example) of trials associated with different awareness ratings which would otherwise be removed by baseline correction (the baseline correction procedure results in each trial’s pre-stimulus window having a mean activity of zero across all time points for each channel, such that our RSA and decoding analyses would be unable to detect and characterise any pre-stimulus contribution to visibility codes).

#### Representational similarity analysis

Representational similarity analysis (RSA) allows us to directly compare bespoke hypotheses about the structure of neural data (Kriegeskorte & Kievit, 2013). In RSA, hypotheses are expressed as model representational dissimilarity matrices (RDMs), which define the predicted similarity of neural patterns between different conditions according to each hypothesis. In our case, we defined 4 model RDMs that make different predictions about whether or not awareness ratings generalise over perceptual content, and whether or not each rating leads to a graded activation pattern partially shared by neighbouring ratings (Figure 3A).

**Figure 3.**
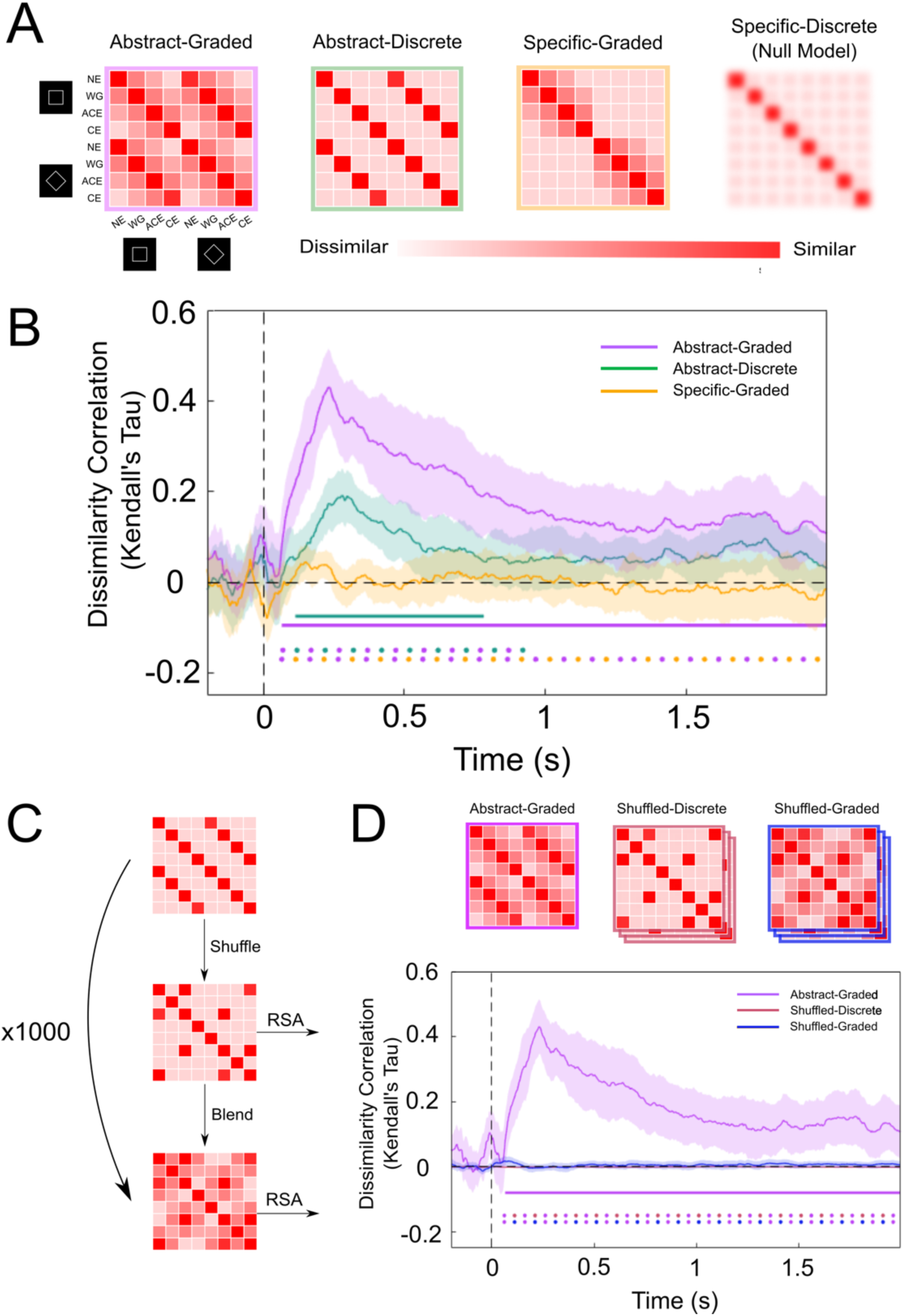
Neural representations of perceptual visibility are abstract and graded. **A***: From left to right:* Abstract-Graded model where neural correlates of awareness ratings are independent of perceptual content and follow a graded structure; Abstract-Independent model where awareness ratings are independent of perceptual content but do not follow a graded structure; Specific-Graded model where awareness ratings are specific to the perceptual content to which they relate and follow a graded structure; Specific-Discrete (Null hypothesis) model where there is no observable representational structure amongst awareness ratings (PAS ratings, NE: No Experience, WG: Weak Glimpse, ACE: Almost Clear Experience, CE: Clear Experience). **B**: RSA reveals that the Abstract-Graded model was the best predictor of the representational structure of neural patterns in whole-brain sensor-level MEG data. Solid horizontal lines represent time points significantly different from 0 for a specific RDM at p <.05, corrected for multiple comparisons. Horizontal dots denote statistically significant paired comparisons between the different models at p <.05, corrected for multiple comparisons. We obtained similar findings across occipital (Figure 3S1A) and frontal (Figure 3S1B) sensors separately, as well as in datasets with stimulus contrast level regressed out (Figure 3S2) and without baseline correction (Figure 3S4). We also examined the pattern of classifier mistakes during cross-stimulus decoding, again revealing distance-like effects in perceptual visibility decoding (Figure 3S3). **C:** Shuffling and blending procedure. This analysis was performed to control for naturally occurring low-frequency content in neural data. **D:** Results from both shuffled models reflect the average Kendall’s Tau over 1000 shuffling permutations. Purple, red, and blue lines represent similarity of the Abstract-Graded, shuffled-discrete, and shuffled-graded models respectively with neural data. The shuffled-discrete line varies only slightly from 0 and is thus hard to see. The Abstract-Graded model is the only model under consideration that significantly predicted the neural data.

In our Abstract-Graded RDM, we model awareness ratings as being independent of perceptual content (such that ratings of a clear experience of a square have an identical neural profile to those of a clear experience of a diamond), as well as being graded in nature (exhibiting a distance effect such that ratings of “no experience” are more similar to those of “weak glimpse”, than of “almost clear experience”). In the Specific-Graded RDM, awareness ratings are modelled as being graded in the same way, but they are now represented differently depending on which specific stimulus they are related to. Conversely, the Abstract-Discrete RDM represents PAS ratings as independent of perceptual content but with no graded structure/distance effects (such that the neural code underpinning a report of “no experience” is equally (dis)similar to the neural code reflecting either a “weak glimpse” or a “clear experience”). Finally, the Specific-Discrete RDM reflects our null model, whereby there is no observable representational similarity structure among conditions, such that neural patterns reflecting one specific awareness rating are equally dissimilar to all other awareness ratings.

RSA involves the comparison of the model RDMs with empirical RDMs constructed from neural data. To do this, we first ran a linear regression on the MEG data with dummy coded predictors for each of our eight conditions (Square trials: NE, WG, ACE, CE; Diamond trials: NE, WG, ACE, CE; trial condition coded with a 1, alternative classes coded with a 0). This resulted in coefficient weights for each condition at each time point and sensor, with the weights representing the neural response per condition, averaged over trials. We then computed the Pearson distance between each pair of condition weights over sensors, resulting in an 8 x 8 neural RDM reflecting the similarity of neural patterns across different awareness ratings and stimulus types (Luyckx et al., 2019). Neural RDMs were subsequently smoothed over time via convolution with a 60 ms uniform kernel.

We then compared this neural RDM with our model RDMs. To compare model RDMs with the neural RDM, we correlated the lower triangle of the model and neural RDMs using Kendall’s Tau rank correlation (Nili et al., 2014). We performed this procedure at every time point, resulting in a correlation value at each time point for each model. Importantly, we only correlated the lower triangle of the RDMs, excluding the diagonal to avoid spurious correlations driven by the increased similarity of on-diagonal values compared to off-diagonal values (Ritchie et al., 2017). This precluded us from directly testing our Specific-Discrete model, since it is represented by a uniform RDM, and as such would give identical rank correlation values regardless of the neural RDM it was compared to. However, since this RDM reflects our null model (i.e. that there is no observable representational structure amongst awareness ratings), this model is implicitly compared with the other model RDMs when we examine whether the correlation of these model RDMs with the neural RDM is greater than 0.

One concern with this approach is that the graded hypotheses may win due to the neural data itself being noisy. In other words, if a classifier is unable to cleanly separate distinct ratings, there will be greater overlap between all ratings in the empirical RDM, including those of close neighbours. To ensure we did not obtain spuriously high similarity with the Abstract-Graded model in virtue of this model’s low-frequency content, we performed a shuffling and blending procedure. This procedure involved shuffling the lower triangle of the Abstract-Discrete RDM before apportioning neighbours of the four (shuffled) high correlation cells with graded amounts of correlation. Correlation was blurred most to immediate neighbours, and less to diagonal neighbours, matching the format of the graded RDMs (Figure 3C). We ran this procedure 1000 times per subject, resulting in 1000 Shuffled-Discrete and 1000 Shuffled-Graded RDMs. We compared all shuffled-discrete and shuffled-graded RDMs with subjects’ neural data at each time point. Finally, we took the average correlation value for each time point across all permutations such that we had a Kendall’s Tau value for both the Shuffled-Discrete and Shuffled-Graded RDMs across time per subject. Through this approach, we were able to compare neural data with RDMs that shared no representational similarity with our Abstract-Graded RDM while controlling for differences in variance and frequency profile.

#### Within-subject multivariate decoding analysis

To support and extend conclusions drawn from our RSA analyses, we ran an exploratory analysis using temporal generalisation methods (King & Dehaene, 2014) to identify content-invariant and graded representations of awareness ratings while also investigating the stability of these representations over time. In this procedure, a separate classifier is trained on each time point (from 200 ms pre-stimulus to 2000 ms post-stimulus) and tested on all other time points. This method results in a time-by-time decoding accuracy matrix indicating the extent to which neural representations are stable over time. Above chance decoding at a particular point in the decoding matrix indicates that neural representations present at the training and testing time points are similar, whilst chance decoding indicates the representations have changed.

We ran the above temporal generalisation analysis using both a within-condition and cross-condition decoding procedure. Within-condition decoding involved training and testing the decoder to classify PAS ratings on trials from one stimulus type (either squares or diamonds). In cross-condition decoding, we trained on trials from one stimulus type and tested on trials from the other (e.g. trained on square trials and tested on diamond trials, and vice-versa). In both cases, we used a 5-fold cross validation scheme, with a balanced number of trials per class within each fold. Cross-condition decoding, where a classifier trained to decode multivariate neural patterns in one class of stimuli is tested on an unseen class of stimuli, offers an empirical test of whether the neural patterns associated with each class share a similar neural code across conditions (Albers et al., 2013; Bernardi et al., 2020; Dijkstra et al., 2018). This analysis therefore complements the RDM analysis in being able to test for content-invariant perceptual visibility codes, while also providing information about their stability over time. We performed all multiclass decoding analyses with a multiclass Linear Discriminant Analysis (LDA) decoder using the MVPA-light toolbox (Treder, 2020) with FieldTrip. Each of the four PAS ratings served as classes for the decoder to classify trials into. We used L1-regularisation of the covariance matrix, with the shrinkage parameter calculated automatically using the Ledoit-Wolf formula within each training fold (Ledoit & Wolf, 2004).

It is important to note that cross-validation is not technically necessary during cross-condition decoding because the test data is never seen by the classifier during training, so there is no risk of overfitting. However, we employed a cross-validation scheme for all our decoders so that differences in their performance would not be due to differences in training procedures (e.g. number of trials in the training or test set). Data was smoothed over 7 samples (28ms) and classification analysis was run on individual time points throughout the whole trial to characterise the temporal dynamics of the representations (−200 to 2000 ms post-stimulus).

#### Stimulus decoding

One difficulty with interpreting a content-invariant representation of perceptual visibility is that it may reflect a lack of power to detect content-specific differences between conditions (e.g. square vs. diamond). To control for this possibility, we sought to ensure that the resolution of our data was sufficiently fine-grained to pick up differences in the neural encoding of different stimuli. To do this, we applied a binary decoding procedure using a binary LDA decoder with the same classification parameters as above. In this analysis, the two stimulus types (squares and diamonds) were used as classes for the decoder to classify trials into. For this analysis we grouped trials into low (NE and WG) and high visibility (ACE and CE) trials to ensure sufficient power, performing the decoding analysis separately in each group. Once again, data were smoothed over 7 samples (28ms) and analysed on individual time points throughout the whole trial (−200 to 2000 ms post-stimulus).

#### Statistical Inference

To determine whether our RSA and decoding results were statistically significant, we used cluster-based permutation testing (Maris & Oostenveld, 2007) with 1000 permutations. For RSA, within each permutation we flipped the sign of each ranked correlation value at each time point for each participant and performed a one-sample t-test against 0. Resulting t-values associated with a p-value smaller than 0.05 were used to form clusters across the single time dimension. For each cluster, an associated cluster statistic was computed, the largest of which was stored per permutation to build a group-level null distribution. The cluster statistic computed from our observed data was then compared to this chance distribution to determine statistical significance with an alpha level of 0.05. This procedure controls for the multiple comparisons problem by only performing one comparison at the inference stage and specifically tests the null hypothesis that the observed data are exchangeable with data from the permuted (null) distribution (Maris & Oostenveld, 2007). We used the same cluster-based permutation procedure to compare how well different model RDMs predicted the neural data. In this case, we performed paired-comparisons where ranked correlation values per RDM were randomly swapped within subjects per permutation to build up a group-level null distribution.

For decoding results, we used the same cluster-forming parameters, but this time randomly flipped the sign of individual subjects’ accuracy scores per permutation to build up a group-level null distribution. Additionally, we formed clusters over both time dimensions of the temporal generalisation matrices. We used the same cluster-based permutation procedure to compare performance between cross-condition and within-condition decoders.

It is important to note that our cluster-based permutation testing procedure does not allow for inference as to the exact time points at which neural representations come into existence. This is because the algorithm does not consider individual time points at the statistical inference stage, since at this point it only relies on cluster statistics, which encompass multiple time points (Sassenhagen & Draschkow, 2019). Still, as we are not interested in the precise onset of content-invariant representations of awareness ratings but rather their general temporal profile, this method is sufficient for our purposes.

### fMRI experiment

To help localise representations of perceptual visibility in the brain, we re-analysed a previously collected fMRI dataset (Dijkstra et al., 2021). It is worth noting that, while source-space decoding in MEG is certainly possible (Andersen et al., 2016; Sandberg et al., 2013), fMRI is much better suited to answering this question at a fine spatial scale, especially as we wish to compare the (potentially fine-grained) differences and similarities in regional activity covarying with perceptual content and/or visibility.

#### Participants

Thirty-seven participants took part in the study. Eight participants were excluded from our analyses. One was excluded because they quit the experiment early, and another because they failed to follow task instructions. The final six subjects were excluded because they did not have at least 10 trials in each visibility rating class after our grouping procedure (see *Within-subject multivariate searchlight decoding analysis* below). Twenty-nine subjects were thus included in our final analyses (mean age = 25.35; SD = 6.31).

#### Stimuli

The stimuli used were taken from the POPORO stimulus data set (Kovalenko et al., 2012). The stimuli selected were a rooster, a fish, a watering can, and a football (Figure 2D), and were selected based on familiarity and visual difference to maximise classification performance as well as both accuracy and visibility scores calculated in a pilot experiment run by Dijkstra et al. (2021). The mask was created by randomly scrambling the pixel values of all stimuli combined (Figure 2C).

#### Experimental Design and Statistical Analyses

The experiment consisted of two tasks: a perception task and an imagery task. Each of these tasks were executed in interleaved blocks and were counterbalanced across participants. Our re-analysis used data only from the perception task and thus we omit details of the imagery component of the study. The perception task ran as follows. A stimulus was presented for 17 ms, followed by a backward mask for 400 ms. Participants then indicated whether the stimulus was animate or inanimate and rated the visibility of the stimulus on a scale from 1 (not visible at all) to 4 (perfectly clear). Button response mappings were counterbalanced across trials. The task was made up of visible and invisible trials. The difference between these trials was the length of the interstimulus interval (ISI) between the stimulus and the mask. In the visible trials the ISI was 66 ms, and in the invisible trials the ISI was 0 ms. In the present study, we only analysed data from invisible trials (Figure 2C) because these were associated with the variation in the visibility ratings that we are interested in. Choosing to focus on a single ISI also means that differences in visibility ratings were not driven by differences in stimulus presentation characteristics. There were 184 trials in total, with 46 repetitions per stimulus divided over 4 blocks. More detailed information regarding the study protocol can be found in (Dijkstra et al., 2021).

#### Acquisition

fMRI data were recorded on a Siemens 3T Skyra scanner with a Multiband 6 sequence (TR: 1 s; voxel size: 2 x 2 x 2 mm; TE: 34 ms) and a 32-channel head coil. The tilt of each participant’s field of view was controlled using Siemens AutoAlign Head software, such that each participant had the same tilt relative to their head position. T1-weighted structural images (MPRAGE; voxel size: 1 x 1 x 1 mm; TR: 2.3 s) were also acquired for each participant.

#### Preprocessing

Data were pre-processed using SPM12 (RRID: SCR_007037). Motion correction (realignment) was performed on all functional imaging data before co-registration to the T1 structural scan. The scans were then normalised to MNI space using DARTEL normalisation and smoothed with a 6 mm Gaussian kernel, which has been shown to improve group-level decoding accuracy (Gardumi et al., 2016; Hendriks et al., 2017; Misaki et al., 2013). Slow signal drift was removed using a high pass filter of 128s.

#### General Linear Model

Coefficient weights were estimated per trial with a general linear model that contained a separate regressor for each trial at the onset of the stimulus convolved with the canonical HRF. Alongside nuisance regressors (average WM and CFG signals and motion parameters), the screen onset and button press of both the animacy and visibility responses were included as regressors, as well as a constant value per run to control for changes in mean signal amplitude across runs.

#### Within-subject multivariate searchlight decoding analysis

For decoding our fMRI data, we binarized the visibility ratings into low and high visibility classes. This is because in contrast to the MEG experiment, visibility was not staircased per-participant, leading to a large number of participants failing to have enough trials at each of the four visibility ratings in both animate and inanimate trials. Because training a decoder on such a small number of trials would yield unreliable and noisy results, trials were therefore sorted into low visibility and high visibility classes on a subject-by-subject basis prior to analysis. This was performed as follows. The median visibility rating (from 1 to 4) was extracted from each subject and trials with a lower visibility rating than the median were classed as low visibility trials, and those with visibility ratings equal to or greater than the median were classed as high visibility trials. This procedure allowed us to control for the fact that different subjects had different distributions of visibility ratings, such that the lower 1 and 2 ratings did not always correspond to low visibility trials, and likewise the higher 3 and 4 ratings did not always correspond to high visibility trials. For instance, one subject may have used visibility ratings 2 and 3 in around 50% of trials, rating 4 on the other 50%, and not used rating 1 at all. In this case, we would label ratings 2 and 3 as low visibility, and rating 4 as high visibility.

Trials were next grouped according to whether they contained an animate or inanimate stimulus. For each participant, if there were less than 10 trials in either the low or high visibility class for either the animate or inanimate trials, the participant was removed. This was the case for 6 participants. The mean number of trials per condition following this procedure were as follows (numbers in brackets denote the standard deviation): animate-high visibility: 63.48 (11.34); animate-low visibility: 25.31 (10.38); inanimate-high visibility: 61.10 (12.24); inanimate-low visibility: 28.86 (11.48).

We used an LDA classifier on the beta estimates per trial to decode low and high visibility ratings within and across animate/inanimate stimulus conditions. Cross-condition decoding was performed by training the LDA classifier on low versus high visibility ratings in animate trials and then testing it on low versus high visibility ratings in inanimate trials, and vice versa. Cross-condition decoding was performed with the same logic as in our MEG analysis: if we train a classifier to decode visibility ratings in animate trials and use this classifier to successfully decode visibility ratings in inanimate trials, we can conclude the representations of visibility ratings are similar across different perceptual content. Once again, we also performed within-condition decoding, where the classifier was trained on low versus high ratings in one condition (e.g., animate trials), and tested on trials in the same condition allow a direct comparison of within- and across-condition decoding performance. This comparison allowed us to determine where content-specific representations of perceptual visibility may exist in the brain. Decoding was performed with a 5-fold cross validation scheme using L1 regularisation with a shrinkage parameter of 0.2, and, similar to the MEG analysis, cross-validation was used for both within-condition and cross-condition decoding. Trials were down-sampled prior to decoding, such that there was an equal number of low and high visibility trials in each fold. To ensure that our data were sensitive enough to show content-specific codes, we additionally ran a similar analysis that sought to decode stimulus content (animate or inanimate) rather than visibility. This analysis was similar in structure except the classifier was trained to decode animate vs. inanimate trials rather than visibility level.

Decoding was performed using a searchlight method. Searchlights had a radius of 4 voxels (257 voxels per searchlight). As such, at every searchlight the classifier was trained on 257 features (one beta estimate for each voxel in the searchlight) for each trial in every fold. The searchlights moved through the brain according to the centre voxel, meaning that each voxel was entered into multiple searchlights. After decoding in each searchlight, the accuracy of the classifier was averaged across folds and this value was stored at the centre of the searchlight to produce a brain map of decoding accuracy.

#### Stimulus decoding in fMRI Regions of Interest (ROI)

As in the MEG analysis, we again wished to establish that findings of content-invariant awareness representations were not due to an inability to decode content itself. We tested our ability to decode perceptual content within two ROIs with successful visibility decoding from our searchlight results. To do this, we created two masks, one visual and one frontal, and then selected the 200 voxels within this mask that had the highest mean visibility decoding accuracy averaged across all four decoding directions (within animate; within inanimate; train animate-test inanimate; train inanimate-test animate). For the frontal mask we used a connectivity-based parcellation of the orbitofrontal cingulate cortex that spanned frontal regions with successful visibility decoding (8m and 32d; (Neubert et al., 2015), whereas our visual mask spanned an area with successful visibility decoding in occipital regions (VO1, VO2, PHC1, and PHC2; (Wang et al., 2015). Using the 200 ROI voxels as features, we decoded animate (rooster and fish) versus inanimate (watering can and football) stimuli in low and high visibility trials separately using the same 5-fold cross validation procedure and LDA parameters as above, down-sampling trials prior to decoding to ensure an equal number of animate and inanimate trials in each fold.

#### Group-level statistical inference

Distributions of accuracy values from classification of fMRI data are often non-Gaussian and asymmetric around chance level. This means that parametric statistical comparisons, such as t-tests against chance decoding (50%), are unable to provide valid tests of whether group-level accuracy values are significant (Stelzer et al., 2013). Therefore, to determine where classifiers had performed significantly above chance, we compared mean performance across all participants with a null distribution created by first permuting the class labels 25 times prior to decoding per participant and then using bootstrapping to form a group-level null distribution of 10,000 bootstrapping samples (Stelzer et al., 2013). We did this separately for each decoding direction (within: train and test on animate; train and test on inanimate; cross: train on animate, test on inanimate; train on inanimate, test on animate). To perform statistical inference on an average cross-decoding map created by averaging the two cross-condition decoding directions, this average map was compared to a group-level null distribution formed by averaging the two null distributions created for the two separate maps. To compare within-condition and cross-condition classification performance, a group-level null distribution was formed by taking the difference between cross and within decoding scores throughout the bootstrapping procedure. To control for multiple comparisons in the searchlight analysis, resulting p values were subsequently corrected for multiple comparisons with a false discovery rate of 0.01.

## Results

### Representational structure of perceptual visibility in whole-brain MEG data

We used RSA to test whether perceptual visibility levels (PAS ratings) correlated with MEG activity patterns independently of perceptual content (Abstract RDMs) or together with perceptual content (Specific RDMs). We additionally tested whether neural activity patterns covaried with visibility levels in a graded or discrete manner (Figure 3A). A model instantiating graded and abstract representations of awareness ratings significantly predicted the neural data throughout most of the post-stimulus period (purple line; Figure 3B). In contrast, a model with an abstract but discrete representational structure was able to predict the neural data only in an early phase of the trial between approximately 100 to 500 milliseconds after stimulus onset (green line). Paired comparisons between these two models showed that the Abstract-Graded model was significantly better at predicting the neural data than the Abstract-Discrete model throughout the majority of the trial (purple and green dots). The Specific-Graded model did not significantly predict the neural data at any point during the trial (gold line), and likewise the Abstract-Graded model was found to be significantly better at predicting the neural data than the Specific-Graded model in a direct comparison (purple and yellow dots), indicating that an abstract model of awareness ratings better described their neural representation. To assess the spatial distribution of Abstract-Graded signals across sensors, we repeated the analysis for frontal and occipital sensors separately (following Hu et al., 2018), finding similar results in each case (Figure 3S1). These results indicate that neural correlates of perceptual visibility generalise over perceptual content and exhibit distance effects, indicative of neural populations tuned to specific degrees of visibility with overlapping tuning curves.

To ensure that the neural data did not exhibit spuriously high similarity with the Abstract-Graded model in virtue of its increased variance and reduced frequency when compared to the Abstract-Discrete and Specific models, we performed a shuffling and blending control analysis (Figure 3C). This procedure revealed no significant prediction of the neural data for either the shuffled-discrete or shuffled-graded RDMs (Figure 3D). As such, RDMs with frequency and variance profiles matching that of the Abstract-Graded RDM, but without any relationship with awareness ratings, were not able to significantly predict neural data, in contrast to the Abstract-Graded model that captures the graded and content-invariant structure of awareness ratings. To additionally control for the possible influence of stimulus contrast on our RDM results, we confirmed that similar results were obtained when regressing out the linear component of contrast (Figure 3S2). It is possible that nonlinear or multivariate effects of contrast may have still driven some of our findings, but as the source of variation in visibility is less important for our main question about content-invariance, we did not pursue this further.

To further characterize the graded representational structure of perceptual visibility, we computed confusion matrices between each rating and its neighbours. By plotting the proportion of predictions for each awareness rating made by the multiclass classifier separately for trials of each rating, we can visualise when our decoder makes mistakes, and which PAS ratings it most often confuses (Figure 3S3). These confusion plots confirm the distance effects identified with the RSA model comparison, in which neighbouring PAS ratings are most often confused with each other by the classifier, and more distant ratings less so, suggesting that visibility is represented in a graded, ordinal manner.

Finally, we asked whether our model RDMs could also predict pre-stimulus neural activity. If a graded, abstract structure for perceptual visibility is already evident prior to stimulus presentation, this would be indicative of trial-to-trial fluctuations in attention or arousal contributing to our ability to decode content-invariant visibility signals. Interpreting (a lack of) pre-stimulus decoding from our previous RSA analyses is confounded by the baseline correction procedure applied during pre-processing. To address this issue, we re-ran our analysis on data that had not been baseline corrected. We found that pre-stimulus activity was not captured by any of the candidate RDMs, and that stimulus-triggered responses continued to show the same graded/abstract pattern of results as in our initial analysis (Figure 3S4). Together these results indicate that pre-trial fluctuations in attention and/or arousal are unlikely to drive our results.

### Temporal profile of perceptual visibility codes

Next, we performed a temporal generalisation analysis to further unpack the content-invariant nature of neural signatures of perceptual visibility and to characterize how and whether their patterns change from timepoint to timepoint. Off-diagonal panels in Figure 4 (top right and bottom left) depict temporal generalisation matrices for both directions of cross-condition decoding (top-right: train on squares-test on diamonds; bottom-left: train on diamonds-test on squares). Within these panels, above-chance decoding on the major diagonal indicates that representations of visibility begin to show content-invariance from just after stimulus onset up until the moment of report. Contrasting cross-condition decoding with within-condition decoding resulted in no significant differences in decoding accuracy for either comparison (train on squares, test on diamonds vs. within squares: all p > 0.89; train on diamonds, test on squares vs. within diamonds: all p > 0.4). In other words, we did not find any evidence that there was content-specific visibility information available over and above content-invariant information. Furthermore, the lack of off-diagonal decoding in each temporal generalisation matrix indicates that the format of content-invariant neural signatures of visibility change rapidly over time.

**Figure 4.**
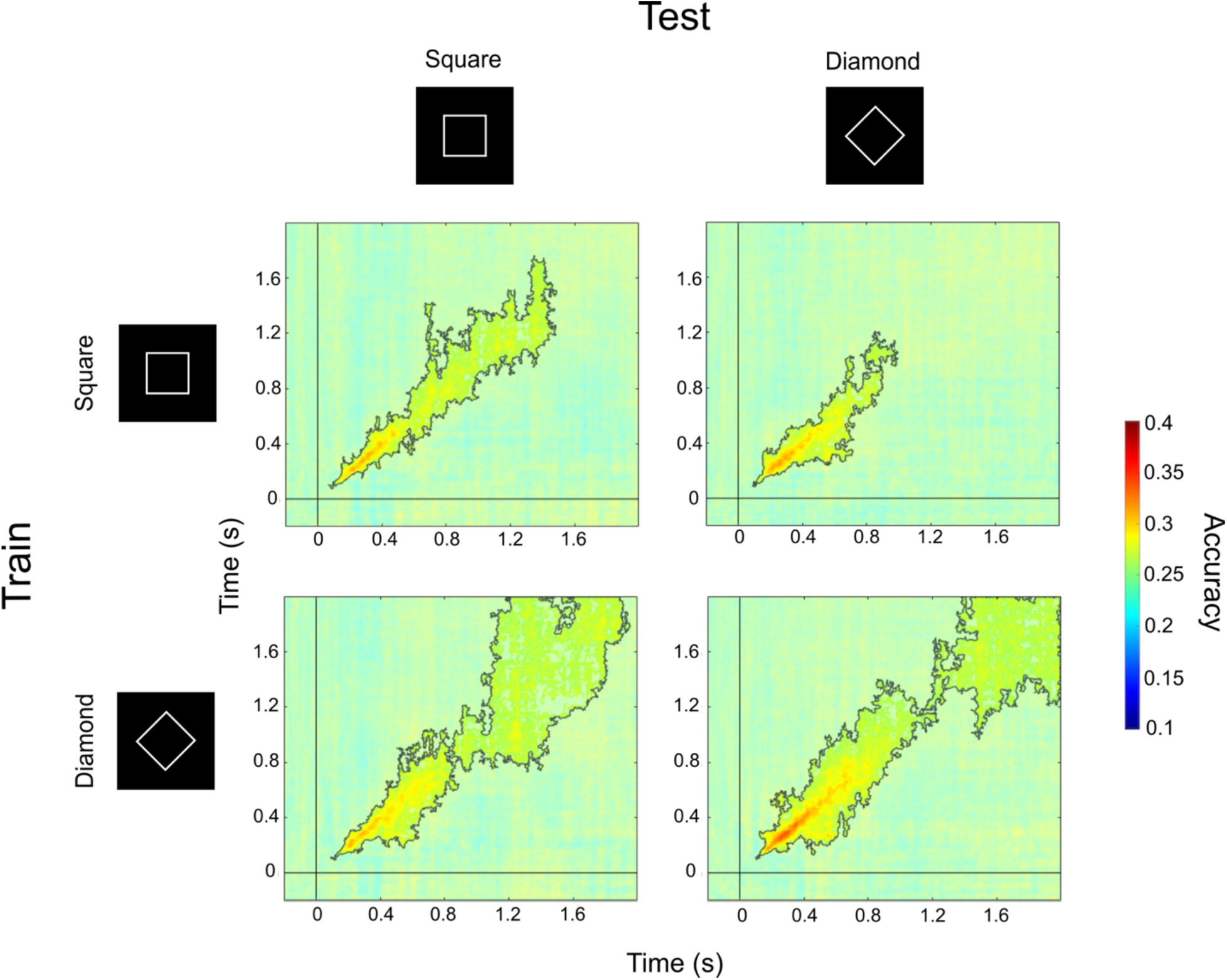
Abstract representations of perceptual visibility evolve rapidly over time. Main figure: Temporal generalisation results for the classification of PAS ratings from MEG data (4 PAS responses; chance = 0.25). For each row, statistical comparisons between the two columns showed no significant differences in decoding accuracy between within- and cross-condition decoding. Non-translucent regions within solid lines highlight above chance decoding, as revealed by cluster-based permutation tests. We replicated these findings in non-baseline-corrected data (Figure 4S1).

We again replicated this analysis in a dataset that had not undergone baseline correction to test whether activity contributing to participants’ awareness ratings could be decoded prior to stimulus presentation. In line with our RSA analysis on this dataset, we could not decode awareness ratings prior to stimulus presentation when data had not been baseline-corrected (Figure 4S1).

### Content-invariant representations of visibility are found across visual, parietal, and frontal cortex

To localise brain regions supporting content-invariant representations of perceptual visibility, we re-analysed an existing fMRI dataset (Dijkstra et al., 2021). We used a searchlight approach to identify brain regions that represent perceptual visibility in an abstract manner. Both cross-condition and within-condition decoding resulted in above chance accuracy in a number of regions across the visual, parietal, and frontal cortex (Figure 5). To assess whether representations of perceptual visibility were stimulus-dependent in certain areas of the brain, we compared cross-condition decoding to within-condition decoding in both animate and inanimate trials (training on animate trials vs. within animate trials; training on inanimate trials vs. within inanimate trials), and found no significant differences. In other words, we found no evidence that stimulus-specific visibility information was present over and above stimulus-invariant visibility information. See Table S1 for details of the clusters of accuracy scores found to be significantly above chance in both cross-condition and within-condition decoding analyses.

**Figure 5.**
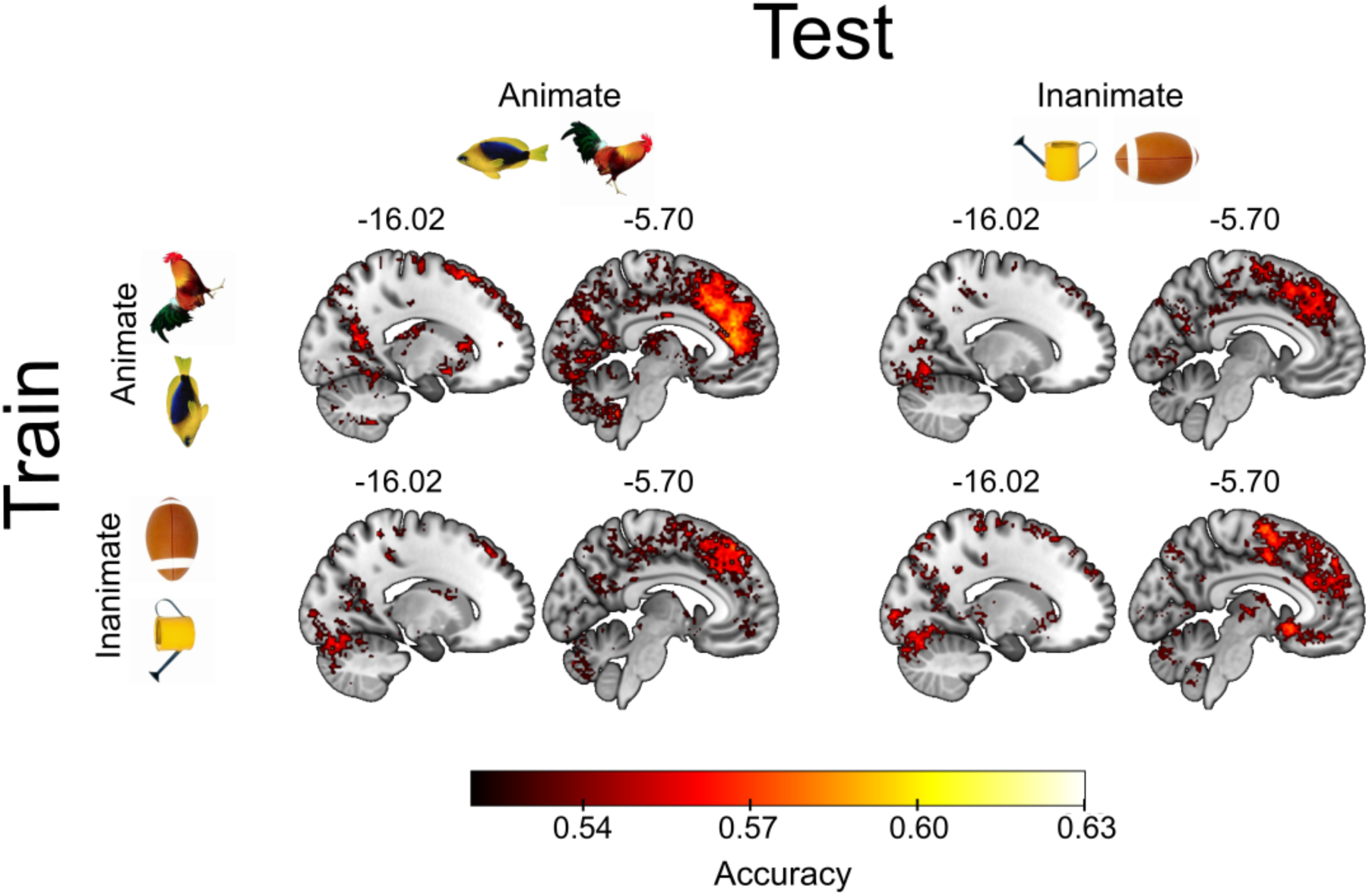
Abstract representations of perceptual visibility are found across visual, parietal, and frontal cortex. Searchlight decoding in fMRI data revealed significantly above chance accuracy in both cross and within-condition decoding of visibility ratings. Clusters of successful decoding were found across large regions of the frontal, parietal, and visual cortex. Our statistical comparison of cross and within-condition decoding accuracy revealed no significant differences anywhere in the brain. Significance was assessed at p < .05, corrected for multiple comparisons with an FDR of 0.01. Clusters are reported in Table S1.

### Stimulus content can be decoded from both MEG and fMRI data

We next considered the possibility that a content-invariant neural signature of visibility may be obtained because of insufficient sensitivity to perceptual content in our dataset. To address this, we sought to decode stimulus identity, rather than visibility level. Stimulus decoding was above chance in both datasets for high visibility trials. In the MEG data, we were able to decode stimulus identity (square vs. diamond) in trials in which participants used the upper two PAS ratings (ACE/CE), but not when participants used the lower two PAS ratings (NE/WG; Figure 6A). Similarly, in the fMRI data, the decoding of animate vs. inanimate stimuli was significantly above chance in a visual cortical ROI during trials reported as high visibility (mean accuracy = 0.52; p = 0.007) but not in trials reported as low visibility (mean accuracy = 0.496; p = 0.655; Figure 6B). It was not possible to decode stimulus content from a frontal cortical ROI in either low (mean accuracy = 0.5; p = 0.406) or high visibility trials (mean accuracy = 0.507; p = 0.159). Together, these analyses indicate that stimulus content could be reliably decoded in posterior visual regions.

**Figure 6.**
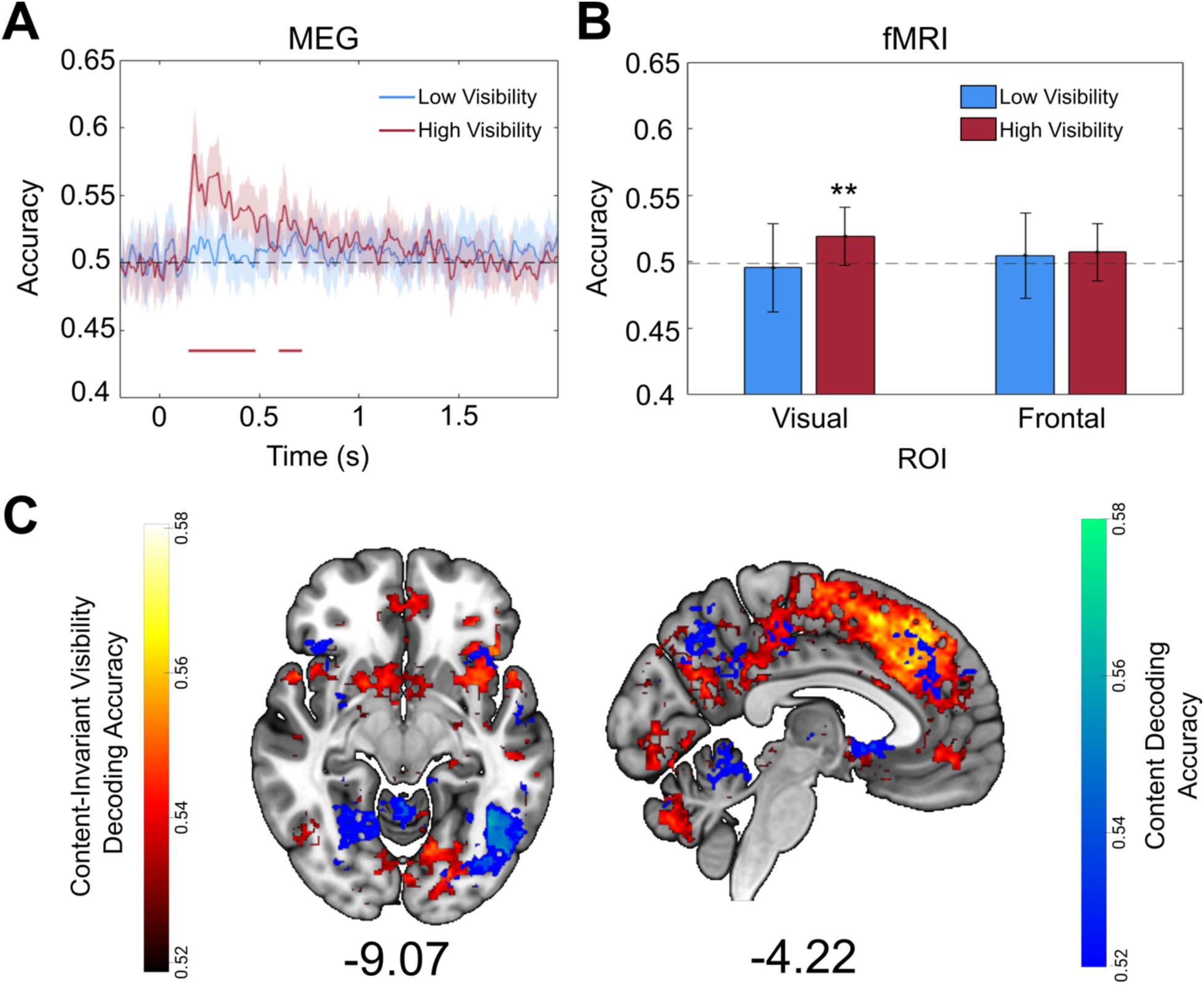
Perceptual content can be decoded in high visibility trials and shows distinct representations to visibility decoding. **A**: Decoding of perceptual content on each trial (squares or diamonds) from participants’ whole-brain sensor-level MEG data for low visibility (NE and WG) and high visibility (ACE and CE) trials separately. Successful decoding was possible in high visibility trials up to approximately 700ms post-stimulus onset. Lines are smoothed using a Gaussian-weighted moving average with a window of 20 ms. Shaded area denotes 95% confidence intervals. The solid horizontal line reflects above-chance decoding, as revealed by cluster-based permutation tests. **B**: Decoding of perceptual content on each trial (animate or inanimate) from participants’ fMRI data for low visibility and high visibility trials separately. Decoding was successful in a visual ROI in high but not low visibility trials, and unsuccessful in a frontal ROI. Asterisks denote significance at p < .01. Error bars illustrate 95% confidence intervals. **C:** Searchlight decoding accuracy for content decoding in high visibility trials (blue) and for content-invariant visibility decoding (red). Clusters illustrate areas where content or content-invariant visibility could be decoded significantly above chance. Content-invariant representations of visibility were more widespread than content representations and extended into the prefrontal cortex, whereas both content and visibility could be decoded in distinct locations of the visual cortex. Significance was assessed at p < .05, corrected for multiple comparisons with an FDR of 0.01. Clusters are reported in Table S2.

### Stimulus Content and Visibility are Encoded in Dissociable Brain Regions

Although perceptual content could be decoded from high visibility trials (Figure 6A, 5B), it is still unclear how these content representations relate to the abstract decoding of stimulus visibility (Figure 5). For instance, it may be that abstract signals reflecting visibility, particularly in the visual cortex, arise from distinct content-specific representations sharing features that covary with visibility.

To further probe the relationship between neural signatures of content and visibility, we ran a searchlight decoding procedure to decode animate versus inanimate stimuli in our fMRI data. Since content could only be decoded in high visibility trials in our ROI analysis (Figure 6B), we restricted the analysis to these trials. We then compared the overlap between the content-decoding searchlight and the content-invariant visibility searchlight maps. To do this we computed mean content cross-decoding accuracy averaged over the two cross-decoding directions (train on animate, test on inanimate; train on inanimate, test on animate) prior to group-level inference (see Methods).

Overall, there was minimal overlap between representations of content and visibility (Figure 6C). Despite overlapping clusters being obtained in the superior and inferior lateral occipital cortex (see Table S2 for full list of individual and overlapping clusters), clear anatomical distinctions in occipital regions can be seen between representations of stimulus content and visibility, with the former being decoded from more lateral regions of the occipital cortex, while the latter was decoded closer to the medial surface (Figure 6C, Table S2). As expected, fewer clusters of above-chance stimulus content decoding were found in frontal regions, whereas content-invariant representations of visibility were more abundant in these areas (Hatamimajoumerd et al., 2022). Distinct decoding patterns for content and visibility representations further strengthens the notion that content-invariant representations of visibility exist partly independently of perceptual content, even in regions typically associated with the encoding of stimulus content such as the visual cortex (Kamitani & Tong, 2005; Kriegeskorte et al., 2008; Mazor et al., 2022).

## Discussion

In this study we asked whether perceptual vividness covaries with neural activity patterns in a content-specific or content-invariant manner. By applying multivariate analyses to MEG and fMRI datasets in which participants rated their awareness of visual stimuli, we found that the vividness of experience is represented in a similar way across different stimulus contents and exhibits signatures of an ordered and graded magnitude code. Furthermore, neural representations of perceptual vividness change rapidly over time and are localized across visual, parietal, and frontal cortices.

The identification of content-invariant representations of perceptual vividness (Figure 3) is in line with recent work highlighting a dissociation between neural correlates of awareness and perceptual content. For example, Sanchez et al. (2020) found neural patterns that indicated whether an individual was aware of a stimulus or not, irrespective of which sensory modality it was presented in. Likewise, Mazor et al. (2022) reported that, while stimulus identity was best decoded from occipital regions, perceptual vividness could be effectively decoded from a wider range of areas including the parietal and frontal cortex. Notably, a recent study also found that graded changes in perceptual vividness could be reliably decoded from the prefrontal cortex, even in the absence of report, consistent with a contribution to the vividness of experience (Hatamimajoumerd et al., 2022). Here we build on these findings by showing that the structure of neural signals underlying graded awareness ratings – ranging from the absence of an experience of particular content, to a clear and vivid experience – are content-invariant.

Our finding that neural representations of awareness ratings display a distance effect (Figure 3; Figure 3S3) is suggestive of perceptual vividness relying on similar schemes to those encoding magnitude in other domains such as number. Specifically, our results are consistent with the possibility that distributed populations of neurons are tuned to specific phenomenal magnitudes, in the same way that specific populations of neurons are sensitive to certain numerical magnitudes (Harvey et al., 2013; Kutter et al., 2018; Piazza et al., 2004). Such a prediction could be tested through repetition suppression experiments (Piazza et al., 2004), and/or by collecting single-unit recordings from human patients while they provide subjective awareness ratings (Pereira et al., 2021). A variety of analogue magnitudes have been shown to rely on common magnitude representations (Luyckx et al., 2019; Pinel et al., 2004; Yallak & Balcı, 2021), prompting a hypothesis that domain-general representations are responsible for encoding low-dimensional quantities in the brain (Summerfield et al., 2020; Walsh, 2003). Therefore, an intriguing possibility is that perceptual vividness is supported by similar domain-general magnitude codes. Future work could explore this hypothesis by assessing whether representations of vividness share neural resources with other analogue magnitude codes such as those for reward or number (Luyckx et al., 2019).

The existence of stimulus-independent representations of perceptual vividness in visual cortical areas (Figure 5) was unexpected, since these areas have been shown to distinguish stimulus features rather than subjective vividness in previous studies (Kamitani & Tong, 2005; Kriegeskorte et al., 2008; Mazor et al., 2022). One concern is that neural representations of vividness ratings as revealed by decoding analyses may look similar across stimuli if content-specific information encoded in separate neural populations is treated as belonging to the same population (i.e. within the same voxel). Successful cross-stimulus decoding of vividness ratings could then occur by way of decoding the amplitude of (content-specific) neural responses in these voxels (Fisch et al., 2009; Moutoussis & Zeki, 2002). As a step towards addressing this concern, we show that stimulus-specific decoding remains possible specifically in visual areas on high- (but not low) visibility trials (Figure 5A; Figure 5B), suggesting that the abstract nature of perceptual vividness representations in this region is not due to a lack of power to detect stimulus-specific effects. Moreover, we show anatomical distinctions between content and visibility encoding (Figure 6C), again indicating that the unexpected above-chance decoding of visibility in the visual cortex is unlikely to be an artefact of a failure to detect content-specific representations.

Another possibility is that content-invariant signals of perceptual vividness in visual cortex reflect pre-stimulus activations that have been shown to contribute to participants’ awareness level in previous studies (Podvalny et al., 2019). Here we could not identify pre-stimulus contributions to visibility codes in our MEG data (Figure 4S2), supporting a hypothesis that the content-invariant and graded representations we report here are largely stimulus-triggered. As such our results suggest that the content-invariant signals related to awareness level in the current data are partly distinct to those reported by Podvalny *et al*. in temporal profile. In any case, it is worth noting that fluctuations in (pre-stimulus) attention and arousal affecting the intensity of experience (as well as other psychological factors like emotional state or motivation) should not necessarily be seen as orthogonal to perceptual vividness signals. Indeed, in certain accounts of consciousness, there is a close relationship between attention-like processes (e.g. the monitoring of the precision or reliability of first-order perceptual content (Feldman & Friston, 2010)), and the phenomenal magnitude attached to that content (Lau, 2019; Fleming, 2020; Graziano, 2013).

By applying temporal generalisation analysis to our MEG data, we were able to reveal the dynamics of vividness representations over time. This analysis indicated that neural patterns covarying with perceptual vividness are unstable, changing during the course of a trial (Figure 4), consistent with a sequence of different neural populations correlating with awareness level over time (King & Dehaene, 2014). Given that we find that vividness is tracked across a variety of cortical regions, such a rapidly changing temporal profile may reflect dynamic message passing between distinct neural populations, consistent with the reverberation of predictions and prediction errors in hierarchical generative models. Future work to directly test this hypothesis could leverage informational connectivity analyses (e.g. Seeliger et al., 2021) to determine the direction of information flow across interacting brain regions, or use RSA to combine M/EEG and fMRI data collected using the same task and stimuli (Cichy et al., 2014).

In summary, we show that perceptual vividness covaries with content-invariant neural representations that exhibit graded distance effects similar to those observed for analogue magnitude codes in other cognitive domains. These representations are spatially distributed and rapidly evolve over time, consistent with the flow of awareness-related information across the visual, parietal, and frontal cortices. This pattern of results adds to growing evidence for a content-invariant neural component contributing to the strength of conscious experience.

## Data Availability Statement

All code is available at the following GitHub Repository: https://github.com/benjybarnett/abstract-awareness. fMRI data is accessible at https://doi.org/10.34973/j9yn-q419 and MEG data will be made available upon acceptance.

## Supplemental Figures

**Figure 2-figure supplement 1.**
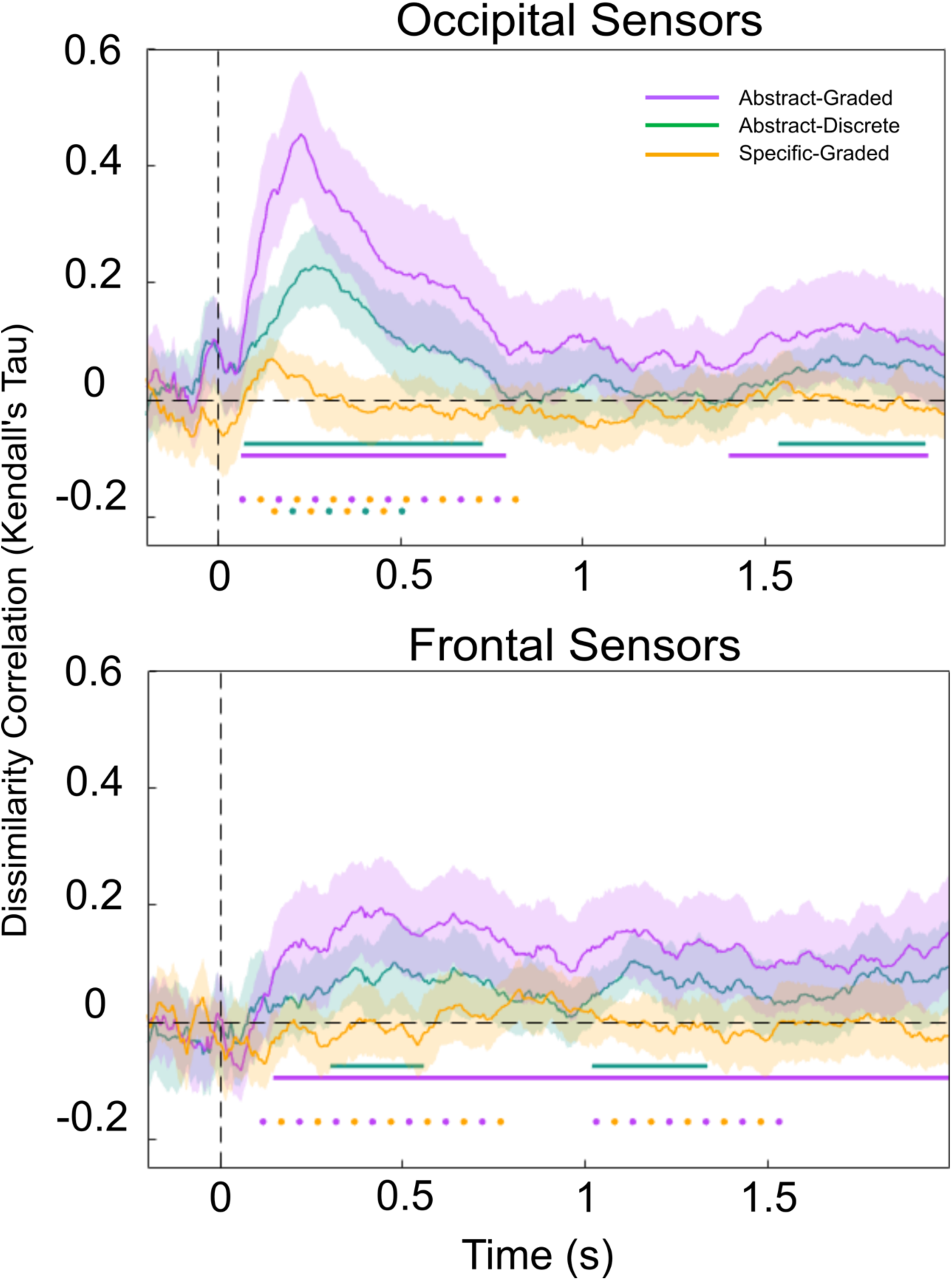
Awareness Ratings Show Similar Representational Structure Across Occipital and Frontal Sensors. RSA analysis performed over occipital sensors (top) and frontal sensors (bottom) only. Purple, green, and gold lines represent similarity of the Abstract-Graded, Abstract-Discrete, and Specific-Graded models respectively with neural data. Solid horizontal lines represent time points significantly different from 0 for a specific RDM at p <.05, corrected for multiple comparisons. The Abstract-Graded model significantly predicted the neural data throughout the majority of the trial (purple line) across frontal sensors, and for a shorter duration when analysis was restricted occipital sensors only. The Abstract-Discrete model was only successful at predicting the neural data across two clusters of time-points post-stimulus when using occipital sensors, but was a significant predictor of neural data for larger portions of the epoch when using frontal sensors. The Specific-Graded model did not significantly predict the neural data at any time point in frontal or occipital sensors. Horizontal dots denote statistically significant paired comparisons between the different models at p <.05, corrected for multiple comparisons. Across frontal sensors, the Abstract-Graded model was a significantly better predictor of the neural data than the Specific-Graded model, and likewise across the occipital sensors, both abstract models significantly outperformed the Specific-Graded model. In this split-sensor analysis, the Abstract-Graded model did not significantly outperform the Abstract-Specific model, however there was a noticeable trend in the same direction as in the RSA performed across all sensors, where the difference in performance was significant (Figure 2B).

**Figure 2 – figure supplement 2.**
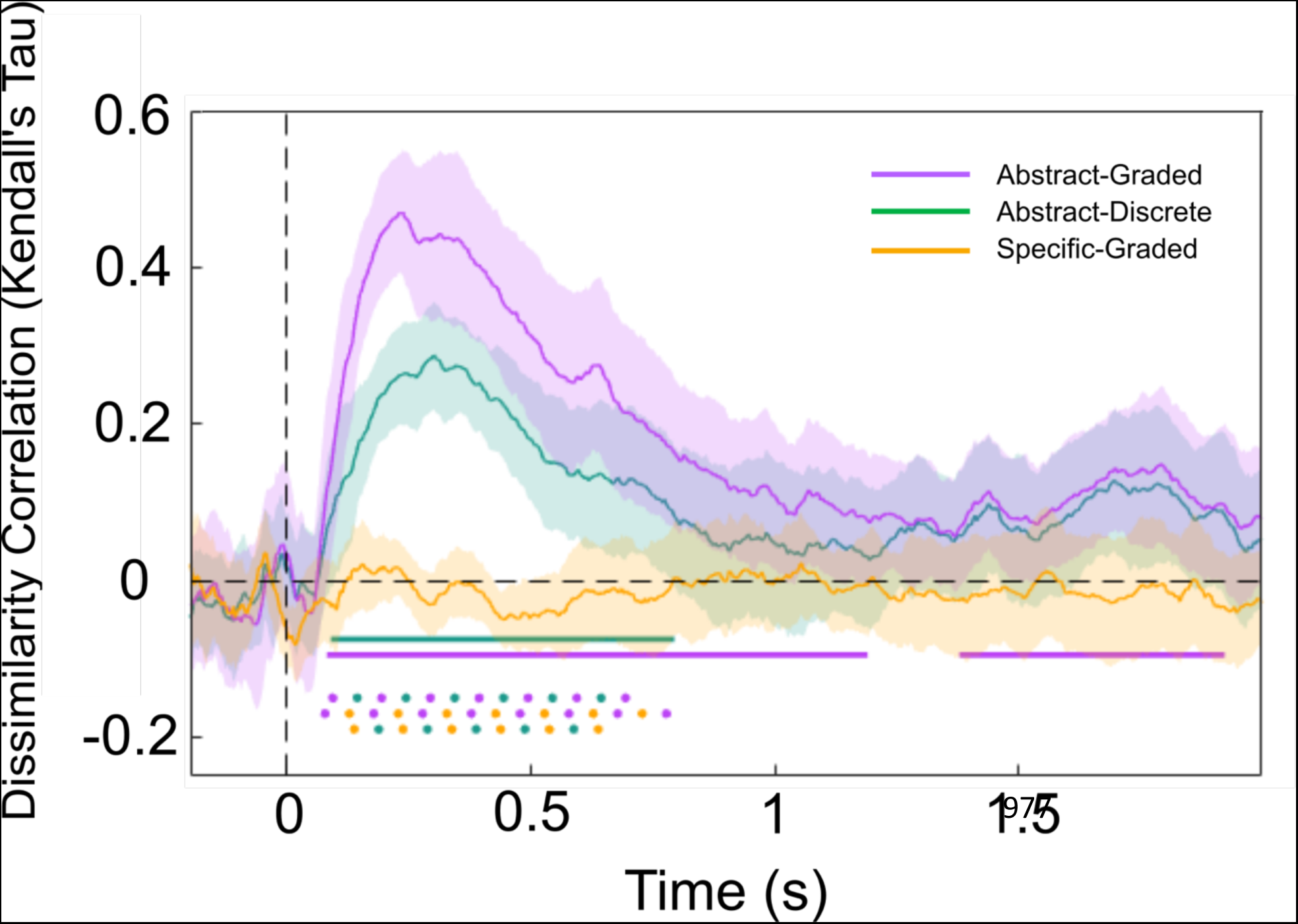
RSA on MEG data with stimulus contrast level regressed out. When stimulus contrast level was regressed out of the MEG data, an RSA still produced comparable results to the original analysis. The Abstract-Graded model still predicted the neural data better than either alternative model. Solid horizontal lines represent time points significantly different from 0 for a specific RDM at p <.05, corrected for multiple comparisons. Horizontal dots denote statistically significant paired comparisons between the different models at p <.05, corrected for multiple comparisons.

**Figure 2 – figure supplement 3.**
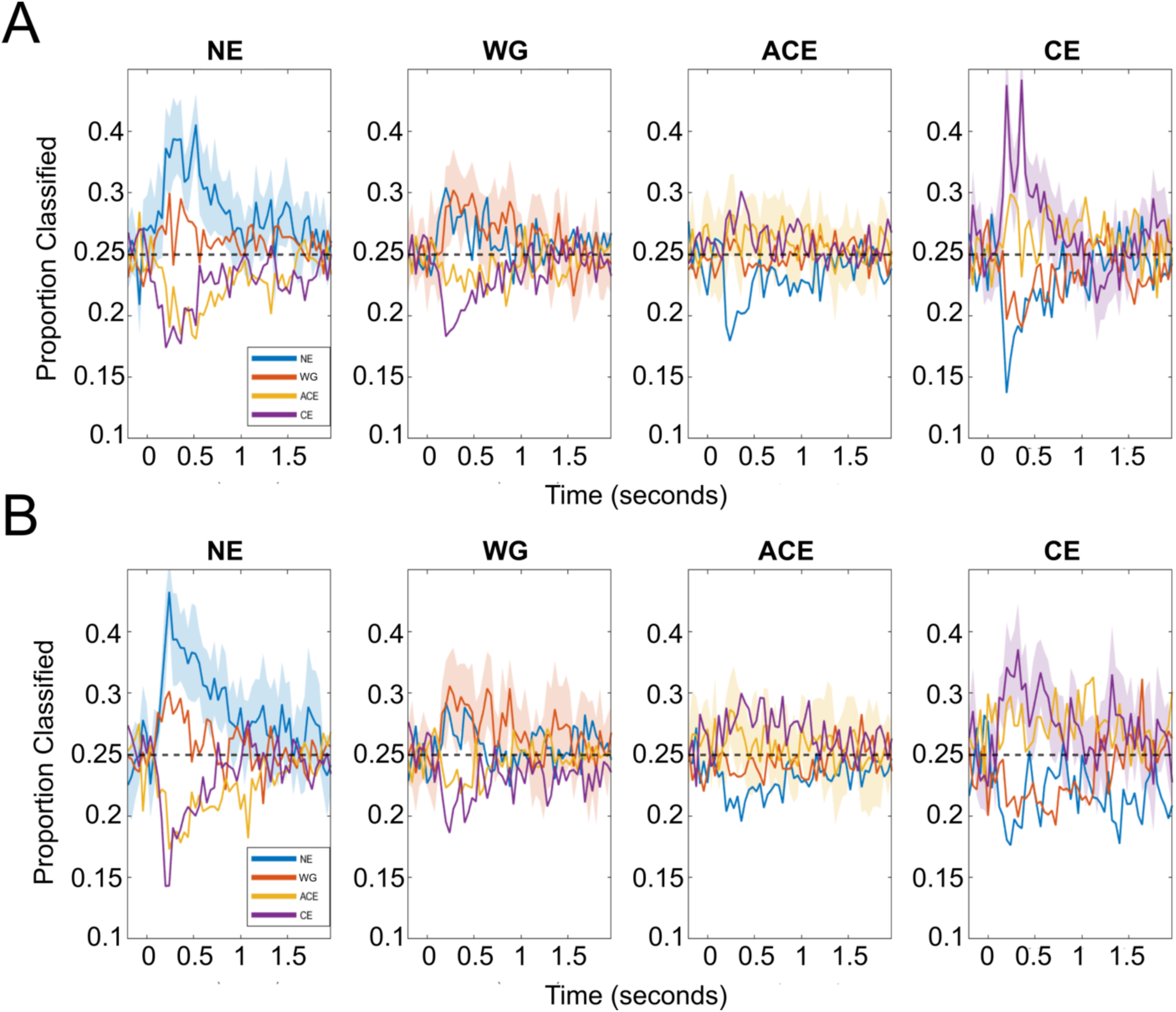
Cross-decoding shows the graded nature of the PAS scale. For each cross-condition decoder (A: Train on Squares; B: Train on Diamonds), the four sub-plots illustrate the proportion of PAS ratings the classifiers decoded trials as. The subplots correspond to trials where the true PAS rating reported by subjects were (from left to right) ‘No Experience’, ‘Weak Glimpse’, ‘Almost Clear Experience’, and ‘Clear Experience’. Each coloured line represents the proportion of trials classified as each PAS rating across time. For example, in trials where participants reported No Experience, the majority of trials were classified correctly (blue line), with the classifier most often misclassifying these ‘No Experience’ trials as ‘Weak Glimpse’ ratings (orange line), and rarely misclassifying them as ‘Almost Clear Experience’ or ‘Clear Experience’ trials (gold and purple lines, respectively). Shaded areas represent 95% confidence intervals.

**Figure 2 – figure supplement 4.**
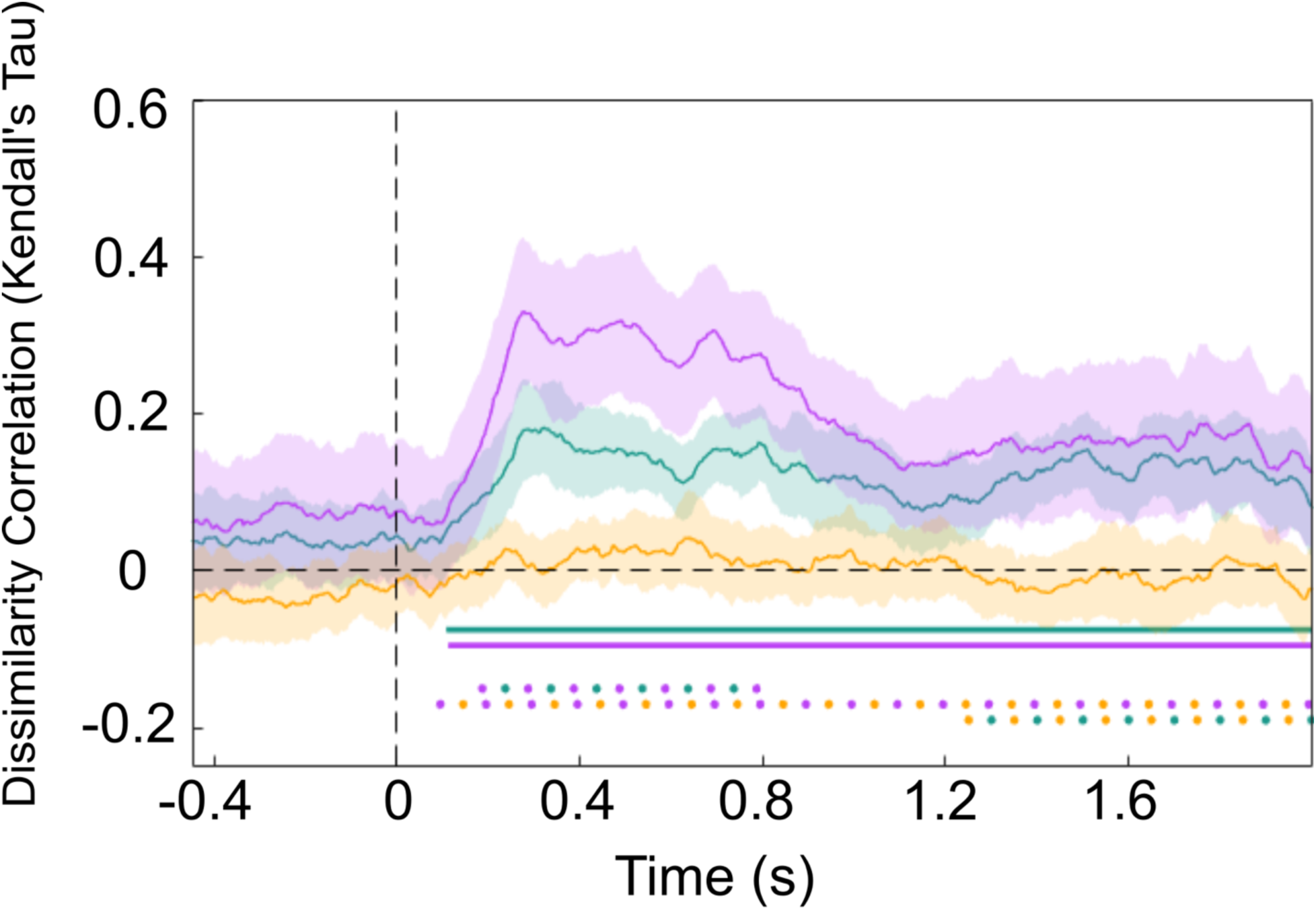
RSA analysis for MEG data without baseline correction. None of our model RDMs significantly predicted pre-stimulus activity in non-baseline-corrected data. Instead, the predominant neural signature was stimulus triggered, as in the main analysis (Figure 2B), with the Abstract-Graded model being the best predictor of neural representations of phenomenal magnitude.

**Figure 3 – figure supplement 2.**
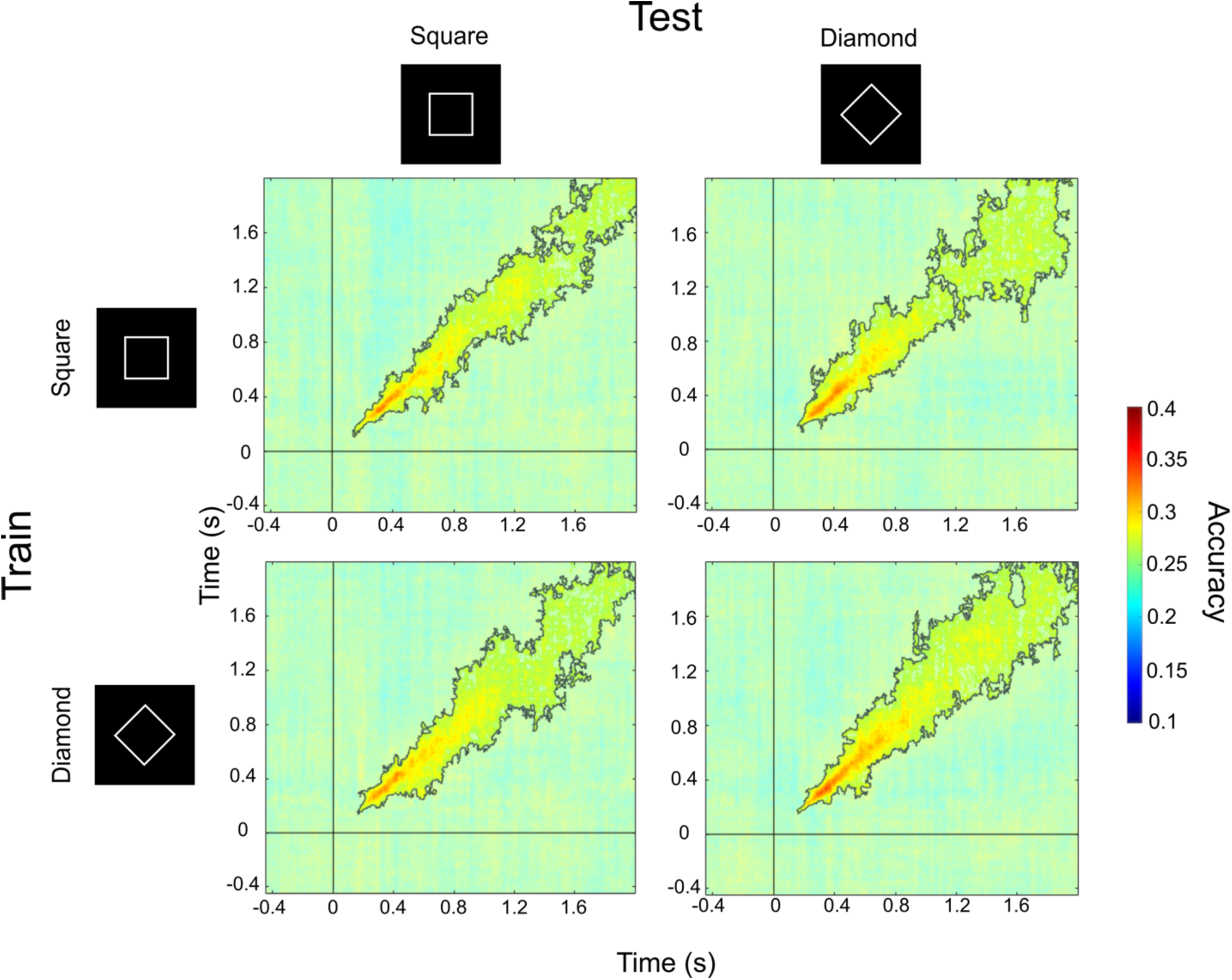
Temporal Generalisation matrices for MEG decoding analyses on data without baseline-correction. For each row, statistical comparisons between the two columns showed no significant differences in decoding accuracy between within and cross-condition decoding. Pre-stimulus decoding of awareness ratings was not possible, even when data had not been baseline-corrected.

**Table S1.** fMRI Searchlight Decoding Results. Clusters with above chance decoding of perceptual visibility for both cross-condition and within-condition decoding. Clusters are significant at p < .05, corrected for multiple comparisons with an FDR of 0.01. Region names are found for the peak co-ordinate using the Harvard-Oxford Cortical and Subcortical Structural Atlas.

| **Decoding Type** | **Atlas Label** | **Cluster Size**<br>**(voxels)** | **MNI**<br>**Coordinates of**<br>**maximum**<br>**accuracy** | **Maximum**<br>**accuracy** |
| --- | --- | --- | --- | --- |
| Cross (train<br>animate) | Postcentral Gyrus | 23,838 | -46, -36, 54 | 0.593 |
| Cross (train<br>animate) | Temporal<br>Fusiform Cortex | 474 | -31.5, -36.1, -25.8 | 0.567 |
| Cross (train<br>animate) | Paracingulate<br>Gyrus | 192 | -8, 44, -6 | 0.563 |
| Cross (train<br>animate) | Frontal Orbital<br>Cortex | 168 | -30, 34, -10 | 0.570 |
| Cross (train<br>animate) | Cingulate Gyrus,<br>posterior division | 109 | 4, -44, 10 | 0.563 |
| Cross (train<br>animate) | Occipital Pole | 89 | -22, -96, 2 | 0.562 |
| Cross (train<br>animate) | Frontal Pole | 75 | 32, 36, -10 | 0.561 |
| Cross (train<br>animate) | Temporal<br>Occipital<br>Fusiform Cortex | 59 | 20, -52, -18 | 0.566 |
| Cross (train<br>animate) | Lingual Gyrus | 56 | 10, -38, -6 | 0.555 |
| Cross (train<br>animate) | Left Cerebral<br>White Matter | 56 | -22, -38, 8 | 0.566 |
| Cross (train<br>inanimate) | Supramarginal<br>Gyrus, anterior<br>division | 23,564 | -54, -36, 30 | 0.593 |
| Cross (train<br>inanimate) | Subcallosal<br>Cortex | 512 | 5.85, 10.6, -6.63 | 0.575 |
| Cross (train<br>inanimate) | Left Caudate | 176 | -10, 2, 16 | 0.575 |
| Cross (train inanimate) | Middle Temporal Gyrus, posterior division | 166 | -52, -42, -2 | 0.585 |
| Cross (train inanimate) | Frontal Orbital Cortex | 152 | -32, 36, -12 | 0.570 |
| Cross (train inanimate) | Left Hippocampus | 106 | -20, -16, -14 | 0.567 |
| Cross (train inanimate) | Central Opercular Cortex | 100 | 38, 0, 18 | 0.571 |
| Cross (train inanimate) | Frontal Pole | 78 | 22, 42, -12 | 0.565 |
| Cross (train inanimate) | Insular Cortex | 71 | -38, -20, -4 | 0.564 |
| Cross (train inanimate) | Lateral Occipital Cortex, superior division | 62 | 32, -88, 14 | 0.555 |
| Within (animate) | Cingulate Gyrus, anterior division | 49,130 | -2, 36, 20 | 0.616 |
| Within (inanimate) | Postcentral Gyrus | 32,710 | 62, -8, 18 | 0.601 |
| Within (inanimate) | Right Caudate | 79 | 18, 10, 16 | 0.565 |

**Table S2.** fMRI Searchlight Content Decoding in High Visibility Trials, Average Cross-Condition Visibility Decoding, and Overlapping Cluster Information. *Decoding type = Content*: Clusters with above chance decoding of perceptual content (animate vs. inanimate) in high visibility trials. *Decoding type = Mean Cross-Condition Visibility*: Clusters with above chance decoding of perceptual visibility for cross-condition decoding when accuracy from both decoding directions (training on animate and training on inanimate) was averaged together. To aid the identification of individual clusters in this map, clustering was performed at an increased accuracy threshold of 0.54. *Decoding Type = Awareness and Content Overlap*: Clusters where decoding of content and cross-decoding of visibility were both successful (i.e. the intersection of the Content and Mean Cross-Condition Visibility clusters). Clusters are significant at p < .05, corrected for multiple comparisons with an FDR of 0.01. Region names are found for the peak co-ordinate using the Harvard-Oxford Cortical and Subcortical Structural Atlas and the MNI Structural Atlas.

| **Decoding Type** | **Atlas Label** | **Cluster Size**<br>**(voxels)** | **MNI**<br>**Coordinates of**<br>**maximum**<br>**accuracy** | **Maximum**<br>**accuracy** |
| --- | --- | --- | --- | --- |
| Content | Inferior Lateral<br>Occipital Cortex | 2768 | -48, -70, -8 | 0.572 |
| Content | Cerebellum | 2150 | 0, -48, -10 | 0.558 |
| Content | Inferior Frontal<br>Gyrus | 1837 | -52, 32, 6 | 0.556 |
| Content | Superior Lateral<br>Occipital Cortex | 919 | -40, - 58, 54 | 0.557 |
| Content | Superior Lateral<br>Occipital Cortex | 907 | -8, -64, 60 | 0.554 |
| Content | Angular Gyrus | 735 | 44, -52, 20 | 0.557 |
| Content | Superior Parietal<br>Lobule | 626 | 20, -44, 62 | 0.548 |
| Content | Frontal Orbital<br>Cortex | 375 | 38, 24, -2 | 0.557 |
| Content | Paracingulate<br>Cyrus | 371 | -6, 26, 34 | 0.547 |
| Content | Inferior Frontal<br>Gyrus | 356 | 46, 20, 20 | 0.563 |
| Content | Superior Frontal<br>Gyrus | 209 | 16, 24, 58 | 0.546 |
| Content | Frontal Pole | 201 | -24, 42, 44 | 0.546 |
| Content | Superior<br>Temporal Gyrus | 192 | -54, -2, -10 | 0.549 |
| Content | Postcentral Gyrus | 189 | -62, -22, 38 | 0.547 |
| Content | Posterior<br>Cingulate Gyrus | 178 | -8, -26, 44 | 0.547 |
| Content | Left Cerebral<br>White Matter | 133 | 6, -24, 16 | 0.544 |
| Content | Cerebellum | 98 | 34, -64, -40 | 0.547 |
| Content | Superior Lateral<br>Occipital Cortex | 76 | 20, -66, 46 | 0.553 |
| Content | Anterior<br>Cingulate Gyrus | 74 | 6, -4, 40 | 0.551 |
| Content | Parietal<br>Operculum Cortex | 56 | 44, -32, 20 | 0.546 |
| Content | Precentral Gyrus | 52 | 44, -16, 58 | 0.542 |
| Mean Cross-Condition<br>Visibility | Precentral Gyrus | 16,638 | -50, 8 32 | 0.584 |
| Mean Cross-Condition<br>Visibility | Postcentral Gyrus | 2863 | -50, -30, 48 | 0.58 |
| Mean Cross-Condition<br>Visibility | Right Accumbens | 278 | 10, 14, -6 | 0.561 |
| Mean Cross-Condition<br>Visibility | Left Cerebral<br>White Matter | 167 | -16, 0, 16 | 0.568 |
| Mean Cross-Condition<br>Visibility | Frontal Medial<br>Cortex | 166 | -4, 48, -10 | 0.555 |
| Mean Cross-Condition<br>Visibility | Middle Temporal<br>Gyrus | 150 | -52, -42, -2 | 0.556 |
| Mean Cross-Condition<br>Visibility | Central Opercular<br>Cortex | 138 | -36, -10, 18 | 0.563 |
| Mean Cross-Condition<br>Visibility | Frontal Orbital<br>Cortex | 133 | 32, 28, 2 | 0.556 |
| Mean Cross-Condition<br>Visibility | Occipital<br>Fusiform Gyrus | 116 | 14, -84, -22 | 0.557 |
| Mean Cross-Condition<br>Visibility | Cerebellum | 103 | -28, -42, -34 | 0.559 |
| Mean Cross-Condition<br>Visibility | Frontal Orbital<br>Cortex | 95 | -36, 36, -10 | 0.563 |
| Mean Cross-Condition<br>Visibility | Temporal<br>Occipital<br>Fusiform Cortex | 91 | -40, -62, -24 | 0.554 |
| Mean Cross-Condition<br>Visibility | Inferior Lateral<br>Occipital Cortex | 58 | 42, -80, 4 | 0.553 |
| Mean Cross-Condition<br>Visibility | Frontal Pole | 58 | 22, 38, -12 | 0.557 |
| Mean Cross-Condition<br>Visibility | Occipital Pole | 56 | 12, -90, -2 | 0.557 |
| Mean Cross-Condition<br>Visibility | Superior<br>Precentral Gyrus | 53 | 26, -12, 52 | 0.552 |
| Visibility and<br>Content Overlap | Cerebellum | 1587 | -6, -82, -32 | N/A |
| Visibility and<br>Content Overlap | Frontal Orbital<br>Cortex | 1521 | -28, 16, -24 | N/A |
| Visibility and<br>Content Overlap | Superior Lateral<br>Occipital Cortex | 782 | -32, -74, 20 | N/A |
| Visibility and<br>Content Overlap | Precuneous<br>Cortex | 474 | -14, -58, 18 | N/A |
| Visibility and<br>Content Overlap | Inferior Lateral<br>Occipital Cortex | 412 | 46, -68, -10 | N/A |
| Visibility and Content Overlap | Supramarginal Gyrus | 383 | 64, -28, 34 | N/A |
| Visibility and Content Overlap | Frontal Pole | 339 | 40, 36, 12 | N/A |
| Visibility and Content Overlap | Cingulate Gyrus | 315 | 0, 36, 16 | N/A |
| Visibility and Content Overlap | Frontal Orbital Cortex | 213 | 36, 22, -14 | N/A |
| Visibility and Content Overlap | Middle Frontal Gyrus | 181 | -24, 36, 32 | N/A |
| Visibility and Content Overlap | Posterior Cingulate Gyrus | 156 | -4, -30, 36 | N/A |
| Visibility and Content Overlap | Middle Frontal Gyrus | 147 | 24, 28, 34 | N/A |
| Visibility and Content Overlap | Right Cerebral Cortex | 145 | 20, 2, -14 | N/A |
| Visibility and Content Overlap | Cerebellum | 115 | 26, -58, -26 | N/A |
| Visibility and Content Overlap | Cerebellum | 111 | -24, -36, -38 | N/A |
| Visibility and Content Overlap | Superior Parietal Lobule | 107 | 34, -50, 36 | N/A |
| Visibility and Content Overlap | Postcentral Gyrus | 101 | -60, -20, 24 | N/A |
| Visibility and Content Overlap | Postcentral Gyrus | 74 | -36, -22, 44 | N/A |
| Visibility and Content Overlap | Lingual Gyrus | 74 | -14, -56, -2 | N/A |
| Visibility and Content Overlap | Cerebellum | 63 | 20, -72, -32 | N/A |
| Visibility and Content Overlap | Cerebellum | 52 | 34, -56, -48 | N/A |

